# Human decellularized extracellular matrix from adipose tissue is a permissive microenvironment for pancreatic organoids generation

**DOI:** 10.64898/2026.03.12.711286

**Authors:** Anastasia Papoz, Sophia Coffy, Florian Jeanneret, Thierno-Sidy Bah, Julia Novion Ducassou, Yohann Couté, Patricia Obeïd, Flora Clément, Christophe Battail, Lisa Martin, Frédérique Mittler, Marta Sacchi, Amandine Pitaval, Xavier Gidrol

## Abstract

*In vitro* reconstruction of human tissue microenvironments that integrate native biochemical and biomechanical cues is essential for disease modelling, regenerative medicine, and personalized therapeutic approaches. However, most currently available engineered matrices fail to recapitulate the complexity and tissue specificity of the human extracellular matrix (ECM).

To address this limitation, we developed a novel hydrogel derived from decellularized human adipose tissue (atdECM) designed to support three-dimensional culture of human cells. The decellularization and delipidation processes were first validated, and the biochemical composition and biomechanical properties of atdECM were comprehensively characterized. Human pancreatic organoids were then cultured within atdECM hydrogel, and their structural organization and transcriptional profiles were analyzed and compared with those obtained in Matrigel, the current gold-standard matrix for organoid culture.

Proteomic and cytokine analyses demonstrated efficient decellularization while preserving collagen-rich ECM architecture and a diverse repertoire of soluble bioactive factors. AtdECM exhibited physiological stiffness and retained tissue-specific extracellular cues. Pancreatic organoids cultured in atdECM displayed morphological similarities with those grown in Matrigel but exhibited transcriptional profiles more consistent with physiological epithelial homeostasis, with reduced activation of inflammatory and stress-related pathways.

Altogether, these findings indicate that atdECM provides a human-derived, tissue-relevant, and permissive microenvironment for human organoid generation. This platform represents a promising alternative to Matrigel for studying human tissue biology and for developing physiologically relevant *in vitro* models.

## INTRODUCTION

Organoids are clusters of cells that are capable of self-organizing and differentiating in a 3D permissive microenvironment. These 3D structures can be derived from embryonic pluripotent stem cells, induced pluripotent stem cells, adult somatic stem cells, primary cell culture from human tissues [1], or even some cell lines (e.g. MCF10A; RWPE1) [2]. With a cell composition close to that observed in the native tissue, organoids partially recapitulate its architecture and function. Thus, they offer an attractive human-based model to study developmental biology, screen pharmaceutical drugs, model diseases, and support regenerative and precision medicine applications [3].

*In vivo*, all human tissues are composed of cells embedded within the extracellular matrix (ECM), a structural scaffold primarily made up of collagens, fibronectin, proteoglycans/glycosaminoglycans, and other glycoproteins. The ECM not only defines tissue architecture but also regulates key cellular functions such as differentiation, adhesion, proliferation, and migration [4]. For organoid development, a physiological 3D environment, including ECM and appropriate conditioned media, is essential. Currently, the most widely used hydrogels for organoid culture are animal-derived, such as Matrigel, which is extracted from Engelbreth-Holm-Swarm murine sarcomas. These hydrogels have significantly contributed to advancements in organoid research [5]. However, their complex composition and tumor-derived origin limit their application in regenerative medicine [6] [7].

To address these challenges, various alternative hydrogels are being developed to better mimic the natural ECM and support organoid formation [8] [9][10] [11]. However, despite these efforts, current systems still fall short in fully reproducing the complexity of the native 3D tissue microenvironment, particularly in replicating tissue-specific mechanical properties and biochemical signals.

As a result, decellularized ECM (dECM) derived from native tissues is emerging as a promising alternative. The decellularization process eliminates nuclear, cellular, and lipid components while preserving ECM proteins, growth factors, cytokines, and non-coding nucleic acids [12]. In this context, dECM-based hydrogels retain much of the native extracellular matrix’s physiological properties, providing essential structural support and signaling cues that promote cell growth and differentiation [13] [14]. This bioactivity, along with the preservation of tissue-specific architecture, offers a physiologically relevant environment for organoid culture and tissue engineering [15].

In this field, animal-derived dECM hydrogels obtained from rat, mice or pig tissues have been proposed to support the development of organoids of liver [16] [17], bile duct [18], small intestine [16] [19], stomach [16] [19], testicular [20], brain [21] and pancreatic islet [21] [14].

Building on these advances, attention has increasingly turned to human tissues as dECM sources. Human-derived ECM more closely mimics the *in vivo* microenvironment and lowers the risk of immunogenicity, making it highly relevant for clinical applications, drug screening, and personalized medicine. Consequently, researchers have expanded decellularization efforts beyond breast [22], liver [23] [24], pancreas [25], peritoneum [26] and intestine [27] to include other human tissues such as kidney [28], lung [29], cardiac/myocardial tissues [30]. All these dECM preparations have been shown to preserve essential biochemical cues and to support cell survival, differentiation and organoid or tissue construct formation *in vitro* and/or *in* vivo. These developments underscore the increasing feasibility of tissue-specific human dECM hydrogels as physiologically relevant scaffolds for organoid culture and translational research.

However, obtaining human tissues for ECM production can be challenging due to limited availability, accessibility, and ethical considerations. Adipose tissue stands out as a notable exception, as it is ubiquitous, routinely discarded during surgical procedures, and available in large quantities, making it an attractive and sustainable source for human dECM production. Importantly, adipose-derived dECM (atdECM) was first pioneered in 2010 by Flynn, who demonstrated that decellularized human adipose tissue provides an inductive microenvironment capable of driving adipogenic differentiation of adipose-derived stem cells [31]. Over the past 15 years, protocols have been refined, optimized and diversified, expanding the applicability of atdECM in tissue engineering and regenerative medicine [32]. Together, these properties (its abundance, accessibility, and biomimetic biochemical composition) make atdECM a particularly promising candidate for providing a supportive and physiologically relevant microenvironment for organoid growth.

In this study, we report the development of a self-gelling hydrogel from decellularized human adipose tissue. We first performed extensive biochemical, proteomic, and biomechanical characterizations of atdECM and compared its properties with those of Matrigel. We then demonstrated that atdECM supports the formation of pancreatic organoids with structural and molecular feature comparable to those obtained in Matrigel. Importantly, transcriptomic analyses revealed that organoids cultures generated in atdECM display gene expression profiles closely aligned with physiological epithelial homeostasis and reduced inflammatory and stress-associated signaling. Together, these findings identify human adipose tissue-derived ECM hydrogels as a robust, accessible, and physiologically relevant alternative to Matrigel, with strong potential for organoid-based disease modelling, drug discovery, and regenerative medicine applications.

## MATERIAL & METHODS

### AtdECM production and characterization

#### Adipose tissue procurement

Fresh human adipose tissues were obtained from Biopredic (Biopredic, France). After reception, samples were immediately stored at -80°C. Tissues were obtained from two female donors, from abdominal sub-cutaneous adipose tissues (see donors characteristics in **Table 1**).

**Table 1:** Donor characteristics.

|  | Donor 1 | Donor 2 |
| --- | --- | --- |
| Age | 66 | 52 |
| BMI | 29 | 28 |

#### Adipose tissue decellularization and delipidation process

Adipose tissues were cut into small pieces (0.5 x 0.5 cm). Samples were rinsed in sterile water for 48 h under agitation. For osmolarity shock, samples were immersed in alternating hypotonic (0.5 M NaCl) and hypertonic (1M NaCl) solutions during two cycles of 2 h each. Following one night of sterile water wash, samples were immersed in 0.05% Trypsin EDTA (1X) (Thermo Fisher Scientific, Waltham, USA) for 1 h 30 min at 37°C to weaken cell membranes. Then, they were homogenized on ice using a Polytron PT 1200 E homogenizer (Kinematica AG, Switzerland) and centrifuged at 900 g for 2 min to favor the phase separation of lipids, which were discarded manually. The protein pellet accumulated at the bottom of the tube was then washed with 20 mL of ultrapure water and immersed in 1% Triton-X100, NH_4_OH 0.1% PBS^-^/^-^ buffer for 72 h at 4°C. Afterwards, samples were rinsed in PBS for 2 days and incubated overnight in 70% isopropanol to remove residual fat and ensure sterilization. Samples were then freeze-dried (Cryotec, Saint-Gély-du-Fesc, France) and ground using a CryoMill (Retsch, Haan, Germany) to obtain a matrix powder that can be stored at -80°C for several months.

#### AtdECM hydrogel

Powders from adipose tissues previously produced were resuspended at 20 mg/ml in pepsin/HCl 0.1 M solution (5 mg/ml) for 4 days under magnetic stirring. The HCl/pepsin activity is neutralized by adjusting the pH at 7.4 with 20 M NaOH solution. Hydrogel solution can be stored several months at -20°C and up to 1 month at 4°C. Before use, hydrogel was sonicated during 2 min at 30 Htz to promote solubilization of large proteins.

#### Longitudinal compression measurement

Pure Matrigel and pure atdECM hydrogels from two donors (donor 1: D1 and donor 2: D2) were cast in a cylindrical mold and left to polymerize at 37°C for 5 h in PBS^-^/^-^. The height of each sample was measured prior to compression testing: Matrigel 3.25 ± 0.27 mm, atdECM D1 2.18 ± 0.52 mm, atdECM D2 1.75 ± 0.27 mm, and fresh adipose tissue 8.67 ± 1.37 mm. To determine the Young’s modulus (E) of the samples, axial compression measurements were performed using a texture analyzer (TA.XT Plus Texture Analyzer, Stable Micro Systems, Surrey, UK). Compression was applied at a speed of 0.1 mm/s, and the force was measured using a 5 kg load cell fitted with a 6 mm-diameter probe. The gels were compressed by 1 mm in the longitudinal direction after reaching a trigger load of 1 g. The compression force (F) was plotted as a function of deformation length (x), and a linear regression was performed over the initial linear (elastic) region. The Young’s modulus (E) was calculated by multiplying the slope of the force-deformation curve by the initial height (H_0_) and dividing by the initial cross-sectional area (A_0_) of the gels:

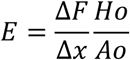

#### Tissue histology

Paraffin section: Fresh adipose tissues and atdECM hydrogels were fixed with 4% formaldehyde and then dehydrated and embedded in paraffin (Leica, Wetzlar, Germany). Slices of 5 µm thickness were cut using a microtome (Leica, Wetzlar, Germany). To assess the effect of the decellularization and delipidation processes, nuclei and collagen were stained using DAPI (0.8 µg/ml) (Roche, Basel, Switzerland) and the PicroSirius kit (Sigma-Aldrich, Saint-Louis, USA) respectively. Images were taken using a Zeiss AxioImager microscope (Zeiss, Oberkochen, Germany) with a color camera (AxioCam105 color) (Zeiss, Oberkochen, Germany).

OCT section: Fresh adipose tissues and atdECM hydrogels were immersed in 2% glutaraldehyde for 1 h at room temperature before being incubated overnight in 30% sucrose at 4°C. Samples were embedded in OCT medium solution (Leica, Wetzlar, Germany) and sectioned at 12 µm using cryomicrotome (Leica, Wetzlar, Germany). To assess delipidation efficiency, slices were labelled with Adipored^®^ (Lonza, Basel, Switzerland). Images were taken using a Zeiss AxioImager microscope (Zeiss, Oberkochen, Germany) with a fluorescent camera (AxioCam MRR3) (Zeiss, Oberkochen, Germany).

#### Mass spectrometry (MS)-based proteomic analyses

Two independent preparations of Matrigel and atdECM hydrogels were prepared. The extracted proteins were solubilized in Laemmli buffer and stacked in the top of a 4-12% NuPAGE gel (Invitrogen, Waltham, USA). After staining with R-250 Coomassie Blue (Biorad, Hercules, USA), proteins were digested in gel using trypsin (modified, sequencing purity, Promega, Madison, USA), as previously described [33]. The resulting peptides were analyzed by online nanoliquid chromatography coupled to MS/MS (Ultimate 3000 RSLCnano and Q-Exactive Plus for atdECM samples and first replicate of Matrigel, and Q-Exactive HF for second replicate of Matrigel, Thermo Fisher Scientific, Bremen, Germany). For this purpose, the peptides were sampled on a precolumn (300 μm x 5 mm PepMap C18, Thermo Scientific, Waltham, USA) and separated in a 75 μm x 250 mm C18 column (Reprosil-Pur 120 C18-AQ, 1.9 μm, Dr. Maisch) using a 120 min acetonitrile gradient. The MS and MS/MS data were acquired using Xcalibur (Thermo Fisher Scientific, Waltham, USA).

Peptides and proteins were identified by Mascot (version 2.8.0, Matrix Science) through concomitant searches against the Uniprot database (*Homo sapiens* taxonomy for atdECM samples or *Mus musculus* taxonomy for Matrigel samples, downloaded on 2022-03-04), and a homemade database containing the sequences of classical contaminant proteins found in proteomic analyses (bovine albumin, keratins, trypsin, etc.). Trypsin/P was chosen as the enzyme and four missed cleavages were allowed. Precursor and fragment mass error tolerances were set at respectively at 10 and 20 ppm, respectively. Peptide modifications allowed during the search were: Carbamidomethyl (C, fixed), Acetyl (Protein N-term, variable) and Oxidation (M and P, variable). The Proline software [34] was used for the compilation, grouping, and filtering of the results (conservation of rank 1 peptides, false discovery rate of peptide-spectrum-match identifications < 1% [35] and minimum of one specific peptide per identified protein group). Proline was then used to perform the compilation, grouping, and MS1 quantification of the identified protein groups. Only protein groups identified with two peptides and quantified with a specific peptide in both replicates of one sample type were considered. For each protein group, intensity-based absolute quantification (iBAQ, [36]) values were calculated from MS1 intensities; the intensities of shared peptides were distributed among the corresponding protein groups based on their iBAQ values calculated using only specific peptides. The iBAQ values of the protein groups in each sample were normalized by the sum of the iBAQ values of all proteins in the sample; the normalized iBAQ values were then averaged for the two analyzed replicates to generate the mean normalized iBAQ value per sample type.

The mass spectrometry proteomics data have been deposited to the ProteomeXchange Consortium via the PRIDE [37] partner repository with the dataset identifier PXD074973.

#### Cytokines & Human Soluble Assays

To compare cytokines and human soluble receptor levels in atdECM hydrogels versus Matrigel (Corning, USA), the Proteome profiler Human XL Cytokine Array Kit, the Proteome Profiler Mouse XL cytokine Array Kit and Proteome Profiler Human Receptor Array, Non-Hematopoietic (respectively ARY022B, ARY028, ARY012) (R&D systems, Minneapolis, USA) were used according to the manufacturer’s instructions. The pixel intensity of each membrane spot was quantified using Image J software (NIH, Bethesda, USA). Signal intensities were background-corrected using blank values and then normalized to the reference spots on each membrane.

#### Atomic Force Microscopy

Atomic force microscopy (AFM) analysis was performed using a Bruker Dimension ICON microscope in TappingMode™ or PeakForce Tapping. The tips used were VTESPA-300 and SAA-HPI-SS, respectively. Error feedback images (“Amplitude error” for TappingMode™ and “PeakForce Error” for PeakForce Tapping) were used to display dECM morphology of each sample.

### Pancreas derived organoids development on atdECM

#### Cell line

Pancreatic H6c7 cells, derived from normal and benign adult human pancreatic tissue via infection with human papillomavirus 16 [38], were purchased from Kerafast company (ECA001-FP) (Kerafast, Boston, USA). Cells were cultured in 2D in Keratinocyte SFM medium (Thermo Fisher Scientific, Waltham, USA) supplemented with 5 ng/ml of human recombinant EGF and 50 µg/ml of Bovine Pituitary Extract (BPE) and used up to passage 21. For passaging, cells were seeded at 25000 cells/cm².

#### Culture of pancreas derived organoids on Matrigel or atdECM coating

##### Matrigel coating

The protein concentration of the Matrigel batch used ranged between 8.8 and 9.1 mg/ml. 40 µl of diluted Matrigel in PBS^-^/^-^ (25%) were deposited in wells of cold 96-well plates (Corning, USA) and directly incubated at 4°C. After 20 min of incubation, the non-gelated Matrigel coating was aspirated.

##### atdECM coating

The atdECM protein concentration ranged between 8 and 9.3 mg/ml. The atdECM hydrogel was briefly centrifuged at 9,000 rpm during 20 sec and diluted at 25% in PBS^-^/^-^. 40 µL of this solution were dispensed in wells of cold 96-well plates and incubated at 37°C to allow gelation. After 45 min of incubation, the ungelled atdECM was removed by vacuum.

##### Cells and top coat deposition

H6C7 cells were trypsinized, counted, and seeded onto Matrigel or atdECM hydrogel coating (400 cells per well of a 96-plate well). For cell sedimentation, the plate was incubated for 45 min at 37°C. Meanwhile, top coats were prepared by adding 4% (v/v) Matrigel in cold Keratinocyte SFM medium supplemented with 5% (v/v) SVF and 5 ng/ml EGF, and gently added on top of the cells. Plates were then incubated (37°C, 5% CO_2_) for 9 days, with media changer every other day.

#### Immunofluorescence of pancreas-derived organoids

To characterize the structures of organoids developed on Matrigel and atdECM coating, organoids were fixed after 9 days of culture with 4% (v/v) formaldehyde (Electron Microscopy Sciences, Hatfield, UK) and 0.8% (v/v) Triton-X100 at 4°C for 25 min, and permeabilized with 0.1% (v/v) Triton-X100 solution for 1h at room temperature. Afterwards, structures were washed three times with PBS Ca^2+^/Mg^2+^ and blocked in 3% (v/v) BSA for 30 min at room temperature. PFA-fixed organoids were incubated with primary antibodies (Table 2) overnight at 4°C and washed three times. Secondary antibodies, including anti-mouse Cy3 (1/500; 115-165-146) (Jackson Immunoresearch, Cambridgeshire, UK) and anti-rabbit Cy5 (1/500; 111-175-144) (Jackson Immunoresearch, Cambridgeshire, UK), were incubated for 5 h at room temperature and washed three times. Nuclei were labelled using DAPI (0.8 µg/ml) (Roche, Basel, Switzerland) for 1 h at room temperature.

**Table 2:**
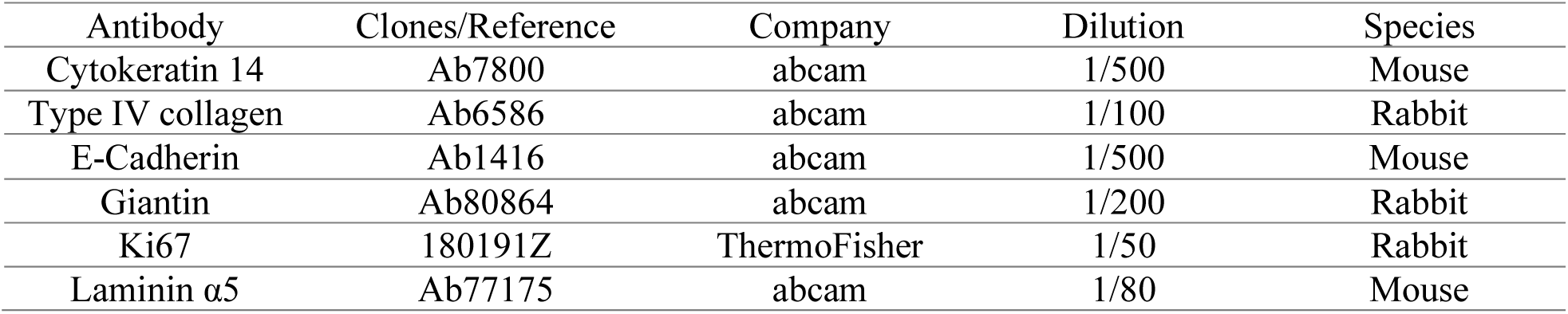
Primary antibodies used to characterize organoids.

#### Image acquisition and quantification

Brightfield images of pancreas derived organoids were taken using a CellInsigth microscope (Thermo Scientific, France). Organoid number quantification was performed manually using ImageJ software (NIH, Bethesda, USA). Immunofluorescence images were acquired using a SpinningDisk confocal microscope (Eclipse TI, Nikon, Japan) and analyzed by MetaMorph and Image J software (NIH, Bethesda, USA).

#### RNA extraction

After 9 days of culture, pancreatic organoids were incubated at 37°C during 2 h in a digestion solution (KSFM media with 1 U/mL dispase (Sigma-Aldrich, Saint-Louis, USA) and 50 U/mL collagenase (Sigma-Aldrich, Saint-Louis, USA)). Once detached from the hydrogel, organoids were resuspended in 0.1% BSA, centrifuged, and further incubated in Accutase (Merk, Darmstadt, Germany) for 1 h 30 min at 37°C. RNA was extracted using RNeasy MicroPlus kit (ThermoFisher Scientific, Waltham, USA) following manufacturer’s guidelines.

#### RNASeq analysis

##### Bulk 3’ RNA-sequencing

Library preparation was performed at the GenomEast platform at the Institute of Genetics and Molecular and Cellular Biology using the Illumina Stranded mRNA Prep Ligation - Reference Guide - PN 1000000124518. RNA-Seq libraries were generated according to the manufacturer’s instructions from 150 ng of total RNA using the Illumina Stranded mRNA Prep, Ligation kit and IDT for Illumina RNA UD Indexes Ligation (Illumina, San Diego, USA). Briefly, oligo(dT) magnetic beads were used to purify and capture the mRNA molecules containing polyA tails. The purified mRNA was then fragmented at 94°C for 8 min and copied into first-strand complementary DNA (cDNA) using reverse transcriptase and random primers. Second-strand cDNA synthesis generated blunt-ended double-stranded cDNA and incorporated dUTP in place of dTTP to achieve strand specificity by quenching the second strand during amplification. Following A-tailing of DNA fragments and ligation of pre-index anchors, PCR amplification was used to add indexes and primer sequences and to enrich DNA libraries (30 sec at 98°C; [10 sec at 98°C, 30 sec at 60°C, 30 sec at 72°C] x 12 cycles; 5 min at 72°C). Surplus PCR primers were removed by purification using SPRIselect beads (Beckman-Coulter, Villepinte, France), and the final libraries were checked for quality and quantified using capillary electrophoresis. Libraries were sequenced on an Illumina NextSeq 2000 sequencer as single-read 50-base reads. Image analysis and base calling were performed using RTA version 2.7.7 and BCL Convert version 3.8.4.

##### Transcriptome bioinformatic analysis

Reads were preprocessed in order to remove adapter, polyA and low-quality sequences (Phred quality score below 20). After this preprocessing, reads shorter than 40 bases were discarded for further analysis. These preprocessing steps were performed using Cutadapt (v1.10). Reads were mapped to rRNA sequences using Bowtie2 (version 2.2.8), and reads mapping to rRNA sequences were removed for further analysis. Reads were mapped onto the hg38 assembly of Homo sapiens genome using STAR (v2.5.3a). Gene expression quantification was performed from uniquely aligned reads using htseq-count (v0.6.1p1), with annotations from Ensembl version 108 and “union” mode. Only reads unambiguously assigned to a gene were retained for further analyses. The ENSEMBL gene IDs were converted into gene symbols using the mapIds function of the R/Bioconductor package AnnotationDbi (v1.60.0). Only genes with a gene symbol and a total sum of reads across all conditions greater than or equal to 10 were kept for further analysis. Gene expression analysis was performed using the R/Bioconductor package DESeq2. Genes with a gene symbol and a total sum of reads across all conditions greater than or equal to 10 were retained for further analysis. Gene expression values were normalized using the variance stabilizing transformation (VST) implemented in DESeq2. Differentially expressed genes (DEGs) between experimental conditions were identified using DESeq2 with the following thresholds: false discovery rate (FDR) adjusted P-value < 0.01 and absolute log2 fold change ≥ 1. The list of DEGs was visualized as heatmaps using the R/Bioconductor package ComplexHeatmap (v.2.14.0) with the following parameters: z-score normalization by row, Pearson correlation as distance method, and use “ward.D2” clustering method. For gene set enrichment analysis (GSEA), genes were ranked using a composite score combining statistical significance and direction of regulation. Specifically, genes were assigned a signed score corresponding to −log10(adjusted P-value), with positive values for upregulated genes and negative values for downregulated genes, based on the sign of the log2 fold change. Hallmark gene sets were retrieved from the Molecular Signatures Database (MSigDB) using the msigdbr R package. Enrichment analysis was performed using the fgsea R package with the following parameters: minimum gene set size = 10 and maximum gene set size = 500. Pathways with an adjusted P-value < 0.05 were considered significantly enriched. Enriched pathways were classified as downregulated or upregulated according to the sign of the normalized enrichment score (NES > 0 or NES < 0, respectively). For each pathway, the leading-edge subset size and the leading-edge ratio (leading-edge genes divided by pathway size) were calculated.

### Statistical analysis

Statistical analyses were performed using GraphPad software (version 7, Graph Pad Software Inc., Los Angeles, CA, USA). Paired Student’s t-tests were applied to compare two variables. Tukey’s multiple comparison test (one-way ANOVA) was applied to compare three or more variables. The p-values are indicated in figure legends. When “ns” or no marks are shown on the graphs, it means that the differences are not significant. Data are represented as mean ± 95% confidence interval [lower 95% – upper 95%]

## RESULTS & DISCUSSION

### Generation and characterization of the adipose tissue-derived ECM

#### Decellularization and delipidation process

To produce dECM derived from adipose tissue, a five-step protocol was implemented, comprising tissue fragmentation, decellularization and delipidation, freeze-drying, cryogenic milling into a fine powder, and enzymatic digestion using pepsin in acidic conditions (**Figure 1A**). The complete procedure required 25 days. Two independent adipose tissue-derived ECM (atdECM) batches were produced from distinct human donors (D1 and D2), using the same protocol. To limit inter-donor variability, adipose tissue samples were obtained from two female donors with comparable BMI values (28 and 29) and from the same anatomical site (abdominal subcutaneous fat, **Table 1**).

**Figure 1:**
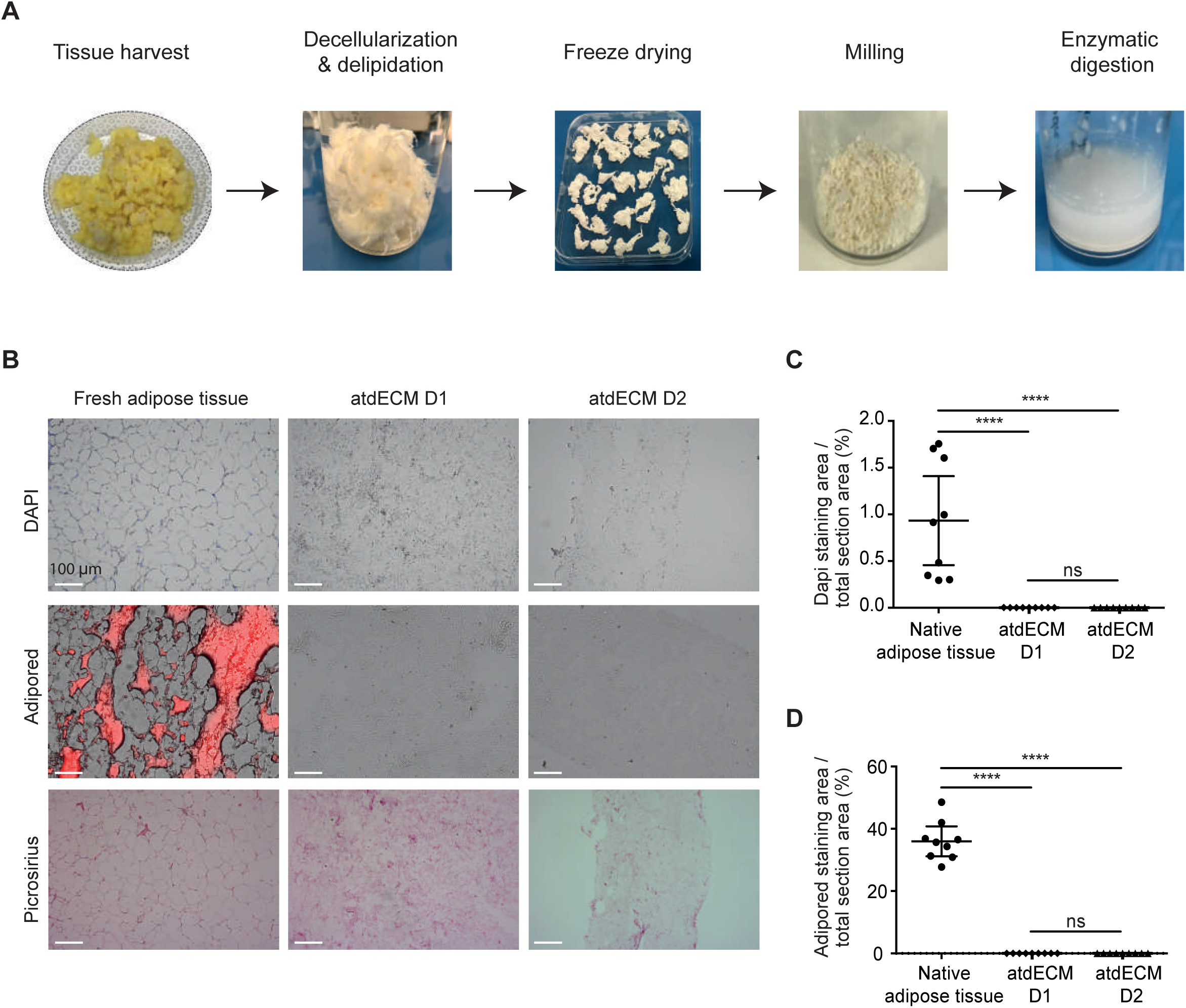
atdECM biological characterization. (A) Adipose tissue decellularization and delipidation process. (B) Sections of fresh adipose tissue or atdECM hydrogel stained by DAPI, Adipored or picrosirius (Scale bars: 100 µm). (C) Quantification of DAPI labeling and (D) Adipored labeling of fresh tissue, atdECM D1 and atdECM D2. (n=3 independent experiments). **** represents p-value < 0.0001.

The efficiency of cell and lipid removal was assessed by histological analysis using DAPI and AdipoRed staining, respectively (**Figure 1B-D**). Compared with native adipose tissue, both nuclear and lipid signals were nearly undetectable in the resulting atdECM hydrogels, indicating efficient decellularization and delipidation. Preservation of ECM structure was evaluated using Picrosirius Red staining, which revealed abundant collagen fibers in atdECM from both donors (**Figure 1B, lower panel**). 100 g of adipose tissue yields approximately 1 g of powder, or 50 mL of hydrogel.

Together, these data demonstrate that the protocol efficiently removes cellular and lipid components while preserving major structural constituents of the ECM. The consistency of the results between donors further supports the robustness and reproducibility of the decellularization and delipidation process.

#### Biomechanical properties of atdECM

The biomechanical properties of dECM matrices are critical determinants of cell behavior, influencing adhesion, proliferation, differentiation, and tissue organization. These properties depend on both technical parameters (e.g., decellularization strategy, protein concentration, pH, and temperature) and biological features such as ECM composition, fibrillar architecture, and tissue of origin [39].

To enable three-dimensional cell culture, atdECM was processed into a hydrogel through lyophilization, pepsin digestion in acidic conditions, and subsequent neutralization. At 4°C, the solubilized atdECM remained liquid, whereas incubation at 37°C resulted in rapid gelation within 1 h, confirming its thermo-responsive self-assembly behavior (**Figure 2A**). This property is essential for homogeneous cell encapsulation and experimental reproducibility.

**Figure 2:**
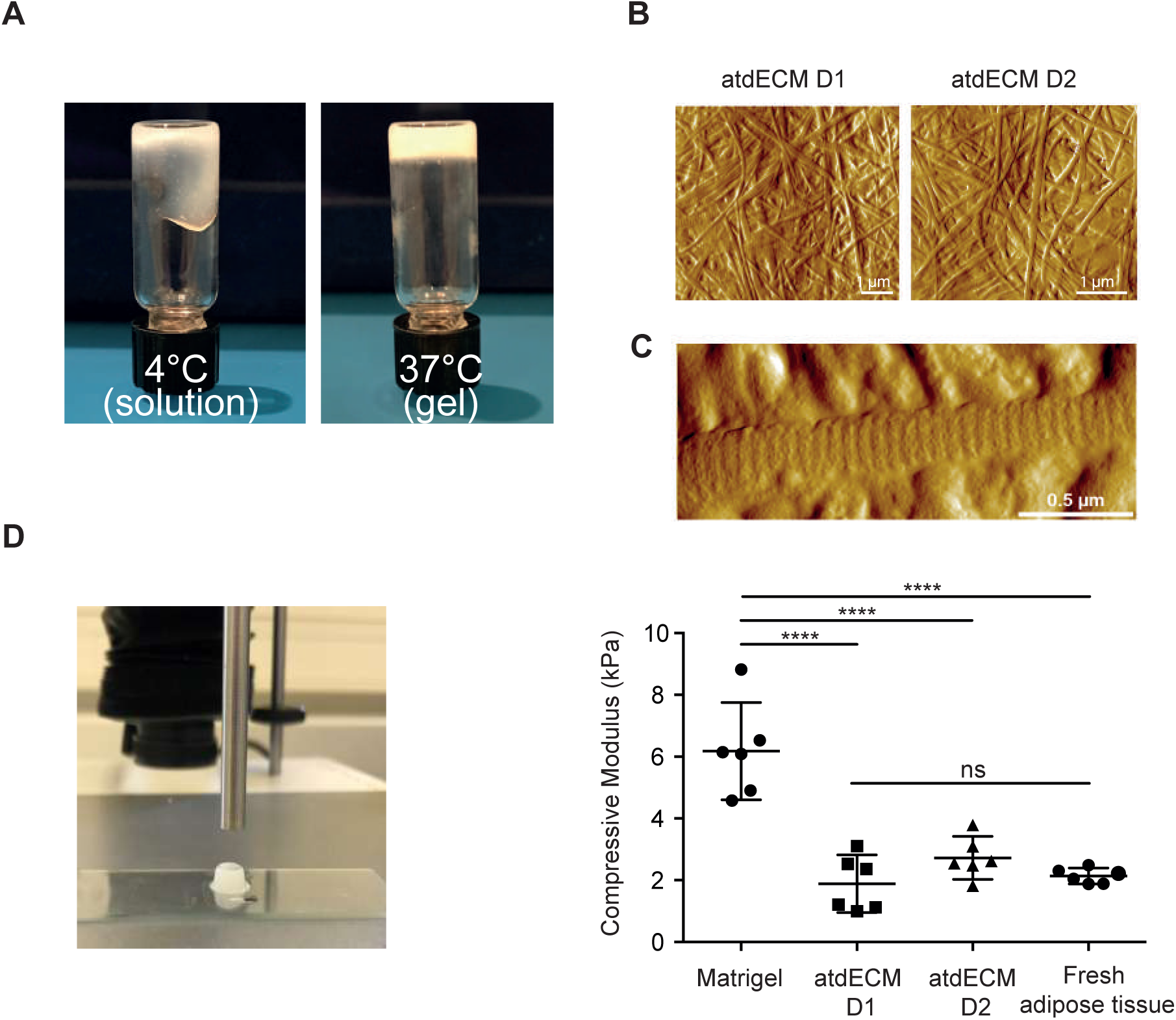
Mechanical characterization of atdECM. (A) Temperature-dependent solution-gel transition of atdECM hydrogel after 1 hour of incubation at 4°C or 37°C (B) AFM images of gelled atdECM D1 and atdECM D2 (scale bars:1µm) (C) Zoom of atdECM D2. Scale bar: 0.5 µm. (D) Picture of the texture analyzer with an atdECM sample. Mechanical properties of Matrigel, atdECM D1, atdECM D2 and fresh human adipose tissue assessed through axial compression tests (n=6 independent experiments). The compressive modulus was used to evaluate matrix stiffness. **** represents p-value < 0.0001.

Atomic Force Microscopy (AFM) was used to further investigate ECM ultrastructure. Interwoven fibrillar networks were observed in all atdECM samples (**Figure 2B-C**), consistent with previous AFM reports identifying such structures as collagen fibers [40]. Notably, the fibers exhibited the characteristic ∼67 nm D-banding periodicity of type I collagen, indicative of preserved native fibril organization, which contribute to both mechanical integrity and cell-matrix interaction.

Mechanical characterization using uniaxial compression revealed that atdECM hydrogels displayed a Young’s modulus of 1.89 ± 0.89 kPa (D1), and 2.73 ± 0.66 kPa (D2), values not significantly different between donors (**Figure 2D**). These moduli closely matched those measured for native human adipose tissue (2.14 ± 0.24 kPa), indicating that atdECM effectively reproduces the physiological elasticity of its tissue of origin. In contrast, Matrigel exhibited a significantly higher compressive modulus (6.18 ± 1.50 kPa), approximately threefold stiffer than both atdECM and native adipose tissue.

Importantly, the Young’s modulus of atdECM (∼2 kPa) is also consistent with reported values for healthy pancreatic tissue prior to tumor-associated desmoplasia [41], and comparable to soft collagen hydrogel commonly used for pancreatic organoid culture [42]. More broadly, these findings align with literature showing that dECM preserves tissue-specific stiffness profiles: soft tissues such as liver [18] or endometrium [43]display low elastic moduli, whereas mechanically constrained tissues (e.g., intestine, colon) exhibit much higher stiffness due to dense fibrillar collagen networks [27].

Preservation of these mechanical cues is likely to influence organoid architecture and cellular organization without imposing artificial lineage bias, supporting the use of atdECM as a permissive and physiologically relevant 3D microenvironment. In summary, atdECM forms a stable thermosensitive hydrogel whose mechanical properties closely resemble native adipose and pancreatic tissues and are markedly softer than Matrigel.

#### Proteomic composition of atdECM

Beyond providing mechanical support, ECMs deliver essential biochemical signals that regulate cell fate and tissue function. Preserving native ECM components during decellularization is therefore critical to recreate physiologically relevant microenvironments.

MS-based proteomic analysis revealed marked differences between atdECM and Matrigel compositions. According to matrisome classification [44], a total of 14 proteins were identified in both atdECM hydrogels, compared with 133 proteins detected in Matrigel (**Figure 3A-B and Figure S1**). atdECM was overwhelmingly collagen-rich, with collagens accounting for 99.95% of matrisome proteins abundances based on iBAQ values. Fibrillar collagens were predominant, with type I representing more than 80% of total collagen and COL3A1 approximately 15% (**Figure 3C and Figure S1**).

**Figure 3:**
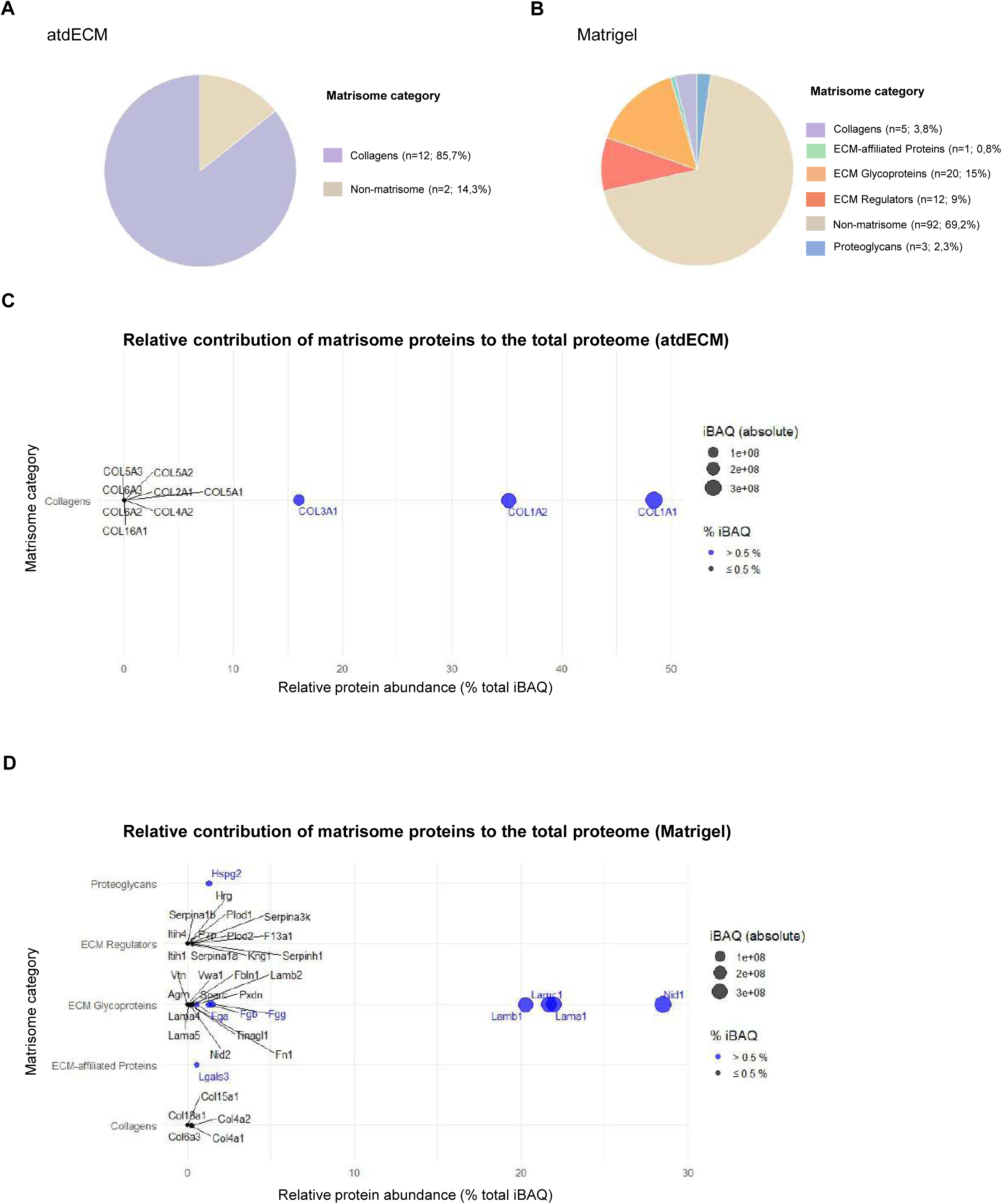
proteomic composition of atdECM Matrigel. (A) Distribution of proteins by matrisome category for atdECM and (B) Matrigel. (C) Dot plot of matrisome protein abundance in atdECM and (D) Matrigel. Relative abundance is expressed as % total IBAQ (x-axis), with dot size proportional to absolute IBAQ and color indicating proteins contributing > 0.5% (blue) or ≤ 0.5% (black).

These findings are consistent with previous reports on atdECM. Using MS-based proteomics, Anderson et al. identified 25 matrisome or matrisome-associated proteins, primarily collagens (29%), along with proteoglycans (29%), ECM glycoproteins (21%), and ECM affiliated-proteins(21%) [45]. Similarly, a more recent study reported collagen type I as the most abundant component across patients regardless of the decellularization protocol used [46]. Moreover, 52 human ECM-specific proteins, including 14 basement membrane-associated peptides, were detected in atdECM, with collagens again representing the dominant class [47]. Notably, our protocol preserves collagen type IV, a key basement membrane component enriched in adipose tissue ECM and implicated in epithelial organization [48]. In contrast, Matrigel exhibited a far more heterogeneous composition, enriched in laminins, nidogens, and other basement membrane proteins, while containing negligible amounts of fibrillar collagen (**Figure 3D**). The limited number of proteins detected in atdECM by MS-based proteomics likely reflects the very wide dynamic range of proteins present in this type of sample, whereby highly abundant collagens can mask lower-abundance proteins for MS detection [49]. This limitation underscores the importance of complementary approaches to detect retained soluble bioactive factors.

Accordingly, cytokine array analyses were performed to profile soluble proteins retained within atdECM hydrogels. AtdECM derived from two independent donors exhibited highly correlated cytokine signatures (Spearman r = 0.9255), indicating strong inter-donor consistency (**Figure 4A**). When compared with previously published datasets using similar, though not identical, array platforms, the atdECM cytokine profile broadly aligned with reports describing human adipose-derived matrices as reservoirs of adipokines, inflammatory mediators, and growth factors, while acknowledging methodological difference that preclude direct quantitative comparisons (**Figure 4B**) [50] [51]. Importantly, our protocol preserved angiopoietin, as well as PDGF and FGF family members, in contrast to the protocol reported by Kim et al’s one [50]. For these factors, our results more closely resembled those reported by Yoo et al. using alginate-fat scaffolds [51]. In addition, our atdECM retained adipose-specific soluble factors such as adiponectin and leptin (**Figure S2A**).

**Figure 4:**
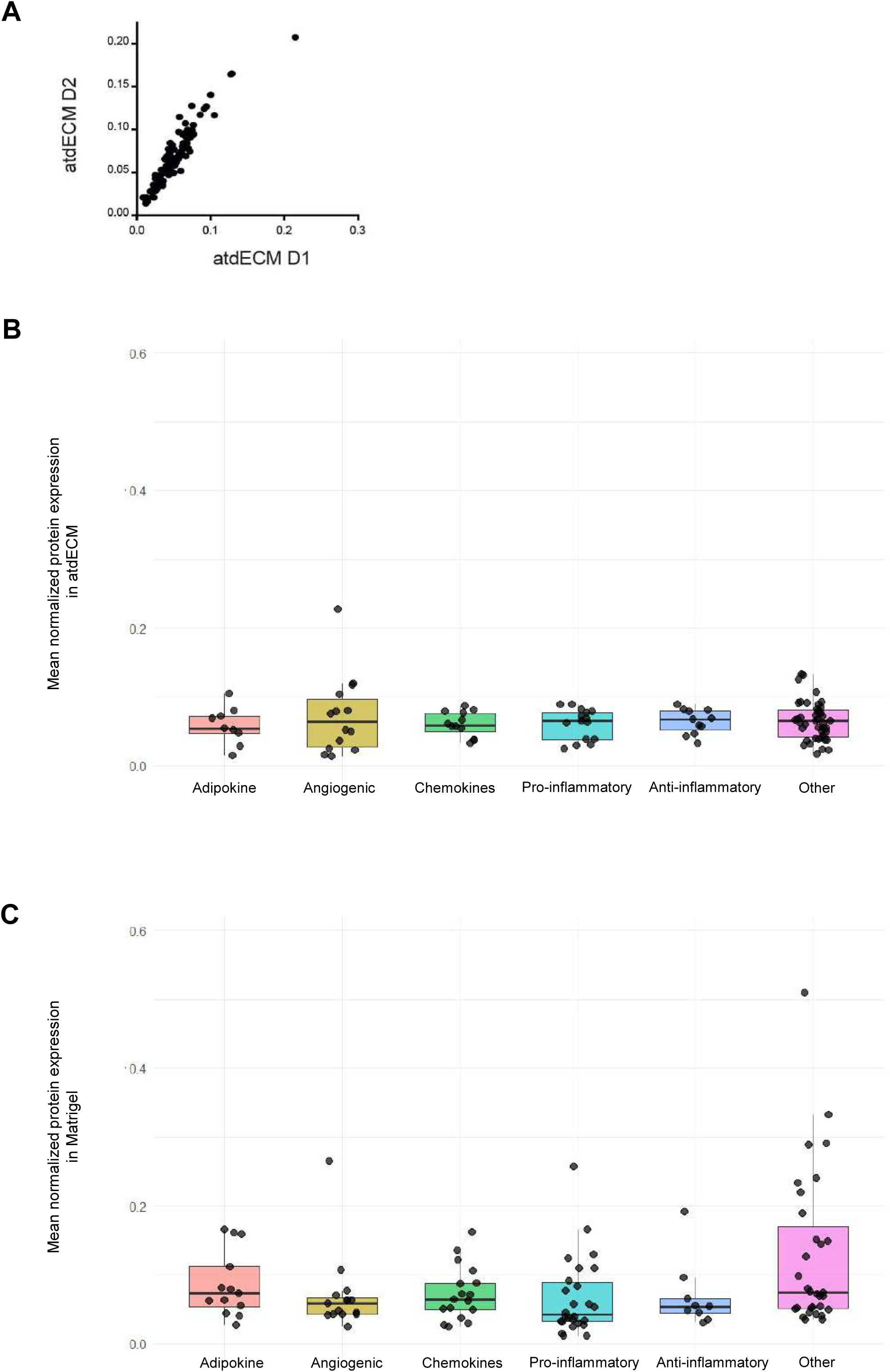
atdECM and Matrigel soluble proteins characterization. (A) Correlation between proteins expressed in atdECM D1 and D2 (R=0.9255). (B) Cytokine expression classified by major cytokine functional categories in atdECM (n=2 donors and 3 membranes / donor) and (C) in Matrigel (n=3).

To position atdECM relative to the reference matrix for organoid culture, cytokine arrays were also performed on Matrigel. Matrigel was strongly enriched in pro-angiogenic and mitogenic factors, including VEGF, Angiopoietin-2, and VCAM-1, as well as matrix remodeling proteins such as MMPs and Serpin E1 (**Figure 4C and Figure S2B**). The Matrigel profile closely matched that reported by Talbot et al., confirming the presence of numerous bioactive growth factors (IGFBPs, VEGF, HGF, FGFs), chemokines (interleukins, CXCLs, CCLs), adipokines (Adiponectin, Leptin, RBP4), proteases, angiogenic and inflammatory mediators (Endoglin, ICAM-1, VCAM-1, Serpin E1), thereby supporting the validity of our measurements (**Figure 4C** and **Figure S2A**) [52]. This intrinsic biochemical complexity reinforces the notion that Matrigel constitutes a highly non-inert and pro-mitotic microenvironment, which may confound the interpretation of organoid phenotypes and highlights the need for more physiologically relevant, human-derived ECM alternatives.

Although direct comparison between human-derived atdECM and murine tumor-derived Matrigel must be interpreted cautiously, relative enrichment analyses revealed clear trends. atdECM displayed a more balanced and tissue-specific biochemical milieu, whereas Matrigel exhibited a strongly mitogenic, pro-apoptotic and pro-inflammatory profile (**Figure 4B-C**), previously associated with elevated inflammatory gene expression in organoids cultured within it [53]. In atdECM, a broader range of cytokines was detected above background, but without dominant outliers (**Figure S2**). Most atdECM-associated proteins exhibited fold changes around two relatives to the median, whereas several Matrigel-associated factors reached fold changes of 5 to 10. This suggests that atdECM provides a more evenly distributed signaling environment, potentially influencing cellular behavior in 3D culture differently from the selectively enriched Matrigel.

Consistent with the literature, dECM retains tissue-specific biochemical fingerprints: soft regenerative tissues (e.g. liver, endometrium) are enriched in collagen types I, III, IV, fibronectin and GAGs, whereas mechanically constrained tissues (e.g. intestine) exhibit denser fibrillar collagen networks and laminins, supporting tissue-specific signalling (**Table 3**). These observations highlight the limitations of generic matrices in reproducing organ-specific biochemical cues. While Matrigel is enriched in laminins, nidogens, growth factors and signaling molecules, it lacks the complexity and tissue-specificity of the native extracellular matrices [54] [55]. In contrast, dECM preserves core matrisome components and maintains the biochemical identity of their tissue of origin [56] [57] [58].

Taken together, proteomic and cytokine profiling demonstrate that atdECM hydrogels differ fundamentally from Matrigel, not only in their structural protein composition but also in the diversity and distribution of soluble bioactive molecules. Whereas Matrigel is characterized by a limited number of highly enriched signaling factors, atdECM displays a broader and more evenly balanced biochemical landscape, which may contribute to the maintenance of physiological cell states in 3D culture.

Although Matrigel remains the gold standard for epithelial organoid culture, its murine tumor origin limits its physiological relevance and raises concerns regarding batch variability and translational applicability, noticeably in regenerative medicine. In contrast, atdECM combines human origin, tissue-specific mechanical properties, and a balanced biochemical composition. These features motivated us to evaluate its capacity to support the generation and maintenance of pancreatic organoids, as explored in the following section.

### AtdECM is a permissive microenvironment for pancreatic organoids generation

#### Phenotypic characterization of pancreas-derived organoids

To assess whether the atdECM hydrogel provides a permissive microenvironment for pancreatic organoid formation, human pancreatic epithelial cells (H6C7) were seeded onto either Matrigel- or atdECM-coated susbtrates and overlaid with a diluted Matrigel layer (2% Matrigel in growth medium) to support epithelial morphogenesis (**Figure 5A**). After 9 days of culture, organoid structures were observed in both coated conditions, whereas no organoid formed in the uncoated control (**Figure 5B**), confirming the requirement for an ECM-derived substrate. Organoids formed on both atdECM and Matrigel matrices displayed exocrine features, organizing into acini-like hollow structures (**Figure 5C**).

**Figure 5:**
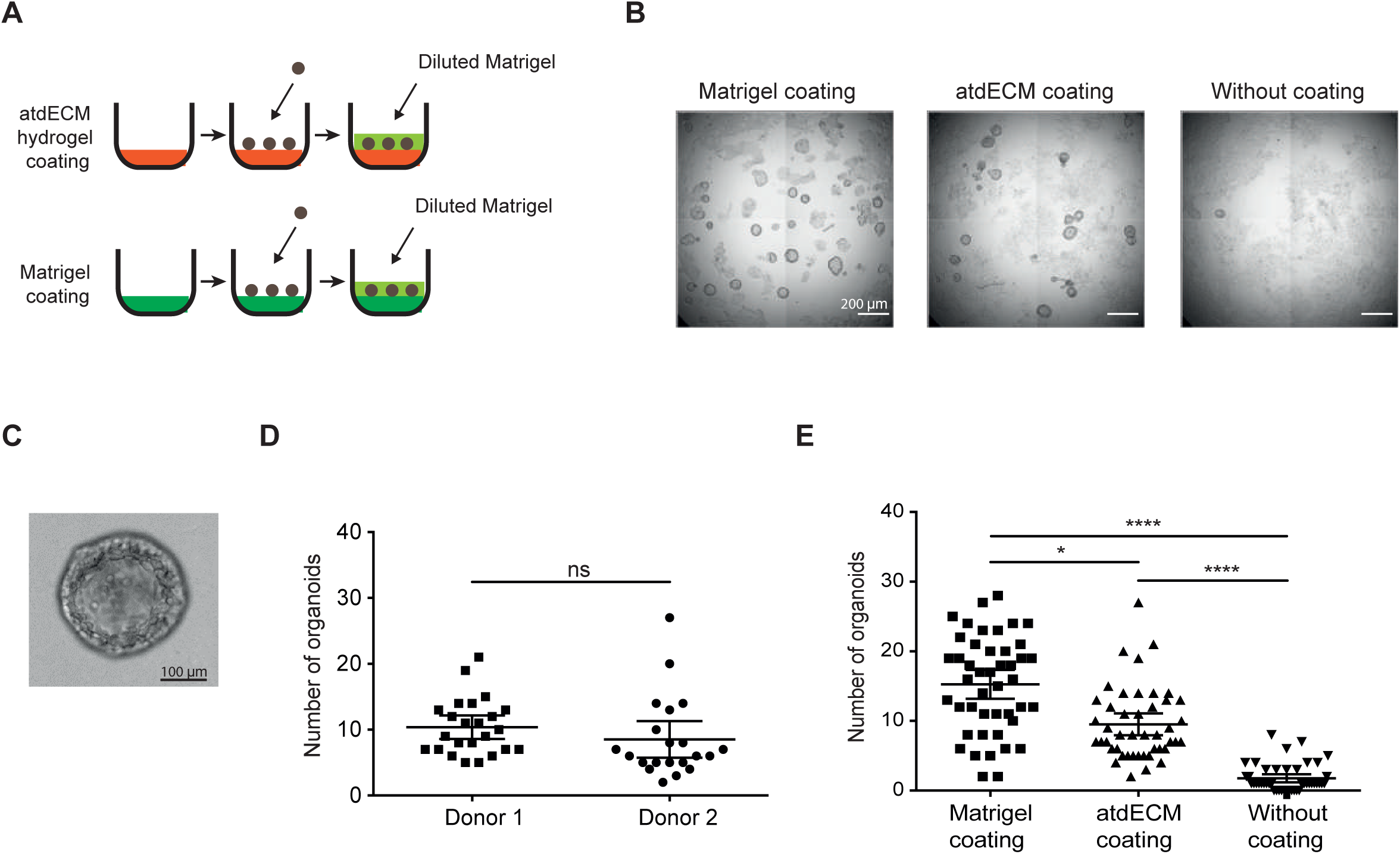
Pancreatic-derived organoids in atdECM coating. (A) Process of pancreatic-derived organoid generation on Matrigel coating or atdECM hydrogel coating. (B) Pancreatic-derived organoids obtained on Matrigel coating (n=2 batches, 3 independent experiments); atdECM coating (n=2 donors, 3 independent experiments); without coating (n=6 independent experiments). Scale bars: 200 µm. (C) Pictures of organoid grown on Matrigel (Scale bar: 100 µm). (D) Number of organoids obtained on atdECM coatings derived from donor D1 and D2 (400 000 cells seeded per well). (E) Quantification of pancreatic-derived organoids generated on Matrigel, atdECM or without coating (n=2 donors, 3 independent experiments) (400 000 cells seeded per well). * and **** represent respectively p-value = 0.0461 and p-value < 0.0001.

Organoid formation was further evaluated using atdECM from two independent donors (D1 and D2). Quantification revealed no significant differences in organoid number between donor-derived hydrogels, with minimal inter-donor variation (**Figure 5D**), demonstrating the robustness and reproducibility of the decellularization process. Based on this consistency, data from both donors were pooled for subsequent analyses.

Counting analysis showed that Matrigel coatings supported a higher number of organoids per well compared with atdECM (15.27 ± [13.19–17.34] *vs*. 9.51 ± [7.95–11.08], respectively). Nevertheless, both hydrogel-coated conditions yielded significantly more organoids than the uncoated control (1.76 ± [1.17–2.34]) (**Figure 5E**).

Organoids cultured on both matrices expressed collagen type IV, cytokeratin 14, and laminin α5, consistent with a basal epithelial phenotype (**Figure 6A-B**). The polarity of organoids seems preserved. Epithelial junctions and Golgi organization were preserved, and Ki67 staining confirmed comparable proliferative activity between conditions (**Figure 6C**).

**Figure 6:**
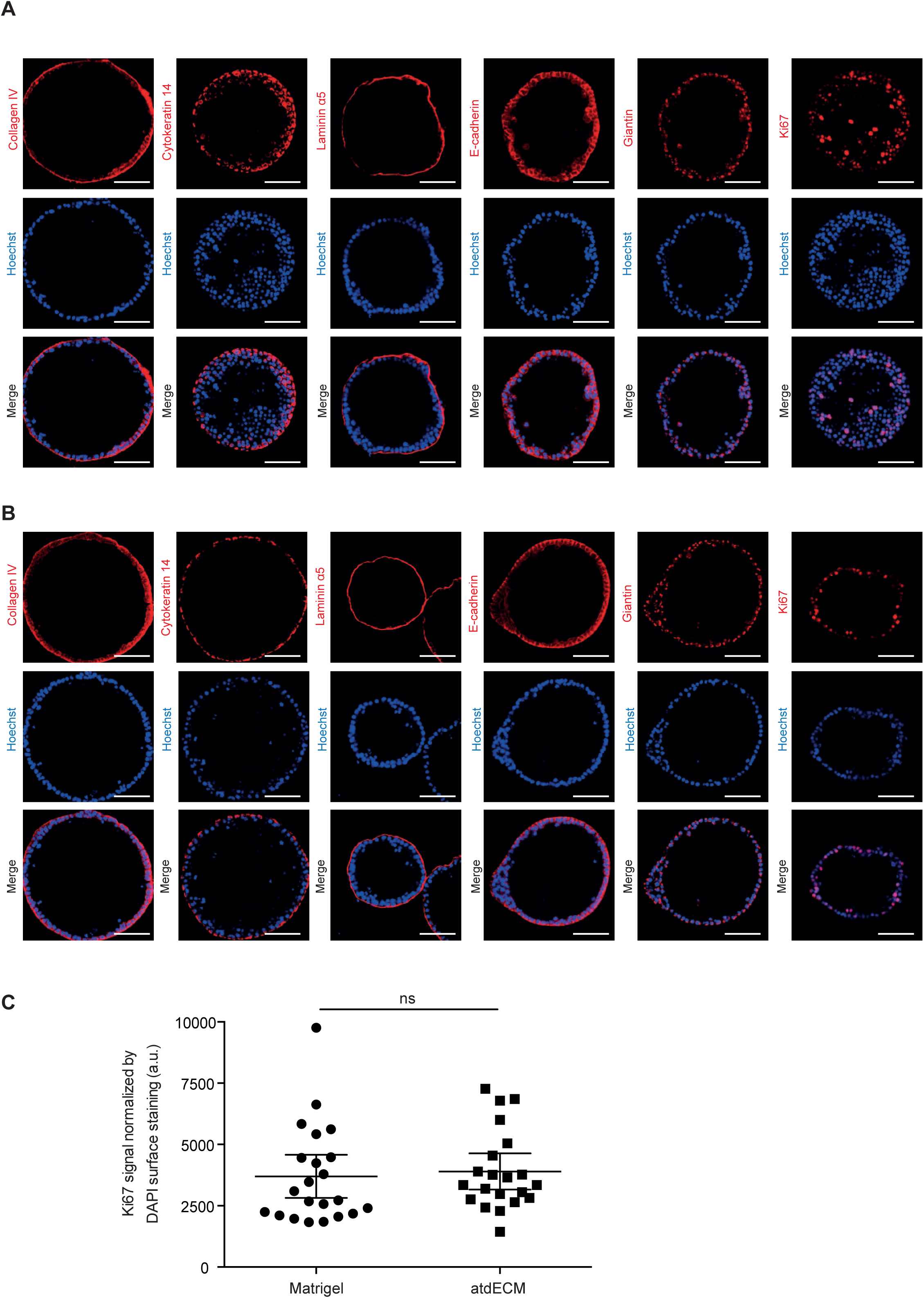
3D culture of pancreatic-organoids in atdECM or Matrigel. (A) H6C7 derived pancreatic organoids in atdECM coating or (B) Matrigel coating stained with Coll IV, Cytokeratin 14, Laminin α5, E-cadherin, Giantin or Ki67. Scale bars: 100 µm. (C) Ki67 signal quantification normalized to DAPI surface staining according to the coating (n=2 donors).

Thus, despite major differences in biochemical and mechanical composition, atdECM supports pancreatic organoids formation with architecture, epithelial identity and proliferative capacity highly comparable to Matrigel. These results show that atdECM provides a microenvironment as permissive qualitatively as Matrigel to generate pancreatic organoids.

#### Transcriptomic profiling of pancreatic organoids

Although organoids cultured on atdECM and Matrigel exhibited similar morphology, we hypothesized that matrix-specific biochemical cues would induce distinct transcriptional programs within organoids. Bulk RNA sequencing was therefore performed to compare pancreatic cell cultures in three conditions: conventional 2D culture, 3D culture on atdECM, and 3D culture on Matrigel.

Gene expression analysis revealed a clear segregation between 2D and 3D cultures (**Figure 7A**), consistent with extensive transcriptional reprogramming induced by 3D culture, largely independent of the specific matrix used [59] [60]. Notably, organoids cultured on atdECM and Matrigel formed distinct clusters, indicating matrix-dependent differences in gene expression despite comparable 3D architecture.

**Figure 7:**
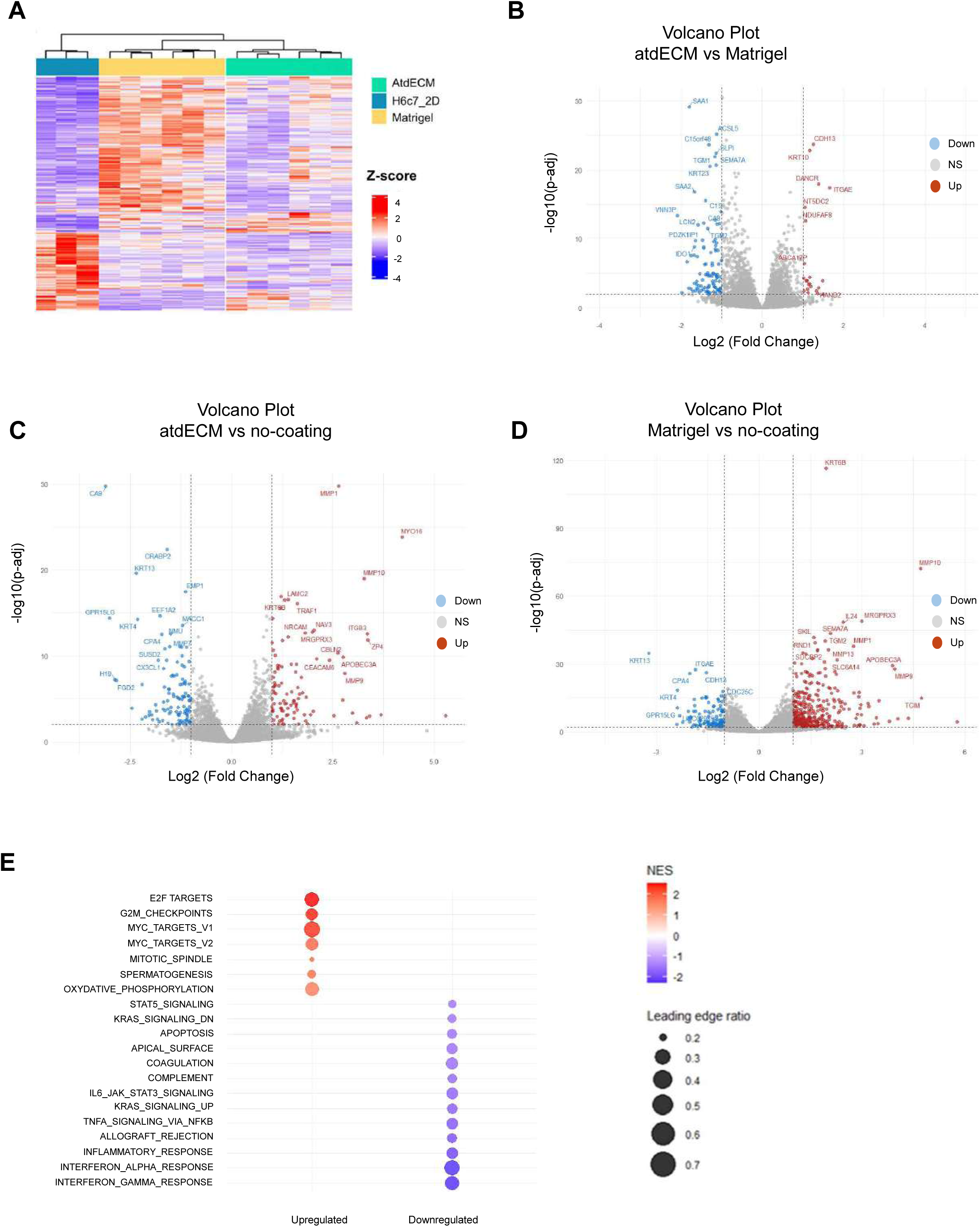
Transcriptomic analysis of pancreatic-derived organoids. (A) Heat map of DEGs identified by pairwise comparisons between atdECM, Matrigel and no-coating conditions using DESeq2 (FDR-adjusted P < 0.01; log2 FC > 1). Expression values are shown as z-scores. (B) Volcano plot of DEGs between organoids cultured on atdECM and Matrigel, (C) between atdECM and no-coating conditions and (D) between Matrigel and no-coating conditions (DEseq2; FDR < 0.01, log2FC > 1). (E) GSEA dot plot of Hallmarks pathways. Color scale represents the normalized enrichment score (NES), indicating activated (positive NES) or suppressed (negative NES) pathways. Dot size reflects the proportion of genes in GSEA leading edge relative to the total pathway size. Pathways are ordered by absolute NES.

Only 122 genes were differentially expressed between organoids grown on atdECM or Matrigel (with 22 upregulated genes in atdECM), compared with 212 and 472 DEGs between atdECM *vs*. 2D culture conditions and Matrigel *vs*. 2D cultures, respectively (**Figure 7B-D**), indicating substantial transcriptional overlap with biologically meaningful differences. The full list of differentially expressed genes (DEGs) is provided in **FigureS3 S4** and **S5**.

Volcano plot analysis revealed significant downregulation of SAA1 and SAA2 in organoids cultured on atdECM compared to those grown on Matrigel (**Figure 7B**). These genes are well-established markers of chronic inflammation and acute-phase responses [61]. Conversely, ITGAE (CD103) and CDH13, genes involved in cell adhesion and intercellular interactions, were upregulated in atdECM cultures [62] [63]. In addition, KRT23, commonly associated with proliferative and stress-associated epithelial states [64], was downregulated, while KRT10, a marker linked to epithelial differentiation, was upregulated [65]. Together, these changes suggest that atdECM promotes a transcriptional program associated with reduced inflammation and enhanced epithelial maturation, consistent with the established role of cell-matrix adhesion in regulating differentiation [66] [67] [68].

Hallmark Gene Set Enrichment Analysis (GSEA) further highlighted pronounced enrichment of interferon-α, interferon-γ, the inflammatory response, and KRAS signaling upregulation pathways in Matrigel-cultured organoids, consistent with its murine tumor origin (**Figure 7E and Figure S7**). These inflammatory pathways were driven by overlapping leading-edge genes, including CD74, RSAD2, CXCL10, RTP4, ISG20, SAMD9L, CXCLC11, GBP4, IFITM2, indicative of a robust interferon-driven inflammatory transcriptional program (**Figure S6**).

In contrast, atdECM-cultured organoids showed enrichment of pathways associated with controlled proliferation and differentiation, including G2M checkpoint, E2F targets, and MYC targets, driven by genes such as PLK1, CDKN2C, CCNA2, CCNB2, and CDC20 (**Figure 7E** and **Figure S6**). Enrichment of oxidative phosphorylation pathways was also observed, consistent with increased metabolic activity associated with controlled proliferation and differentiation. Such metabolic signatures have been linked to stem and progenitor cell fate regulation and tissue homeostasis [69] [70] [71][72] [73].

#### Linking ECM composition to transcriptional outcomes

The transcriptional differences observed between atdECM- and Matrigel-cultured organoids are highly consistent with the biochemical differences identified by proteomic and cytokine analyses. The pro-inflammatory, growth factor-rich composition of Matrigel likely drives interferon and KRAS-associated signaling, as reported in multiple organoid systems where Matrigel induces oncogenic or stress-related gene expression [53].

By contrast, atdECM displayed a more balanced and tissue-specific cytokine profile, characterized by moderate enrichment of adipokines, growth factors, and cytokines, without dominant pro-inflammatory signals. This more evenly distributed biochemical environment is consistent with the reduced inflammatory gene expression and enhanced adhesion- and differentiation-associated transcriptional programs observed in atdECM-cultured organoids. These findings align with previous studies showing that human-derived dECM supports physiological cell states by preserving native biochemical cues [12].

## CONCLUSION

In summary, atdECM supports the generation and maintenance of pancreatic organoids with architecture, epithelial identity, epithelial polarity and proliferative capacity comparable to Matrigel, while inducing a distinct and more physiological transcriptional program characterized by reduced inflammatory signaling and enhanced control of proliferation, differentiation and cell adhesion. These differences are not readily apparent from morphological analyses alone but emerge at the molecular level, highlighting the importance of integrating transcriptomic profiling when evaluating biomaterial performance.

Although Matrigel and atdECM are very different in terms of chemical and physical cues, both allow the generation of morphologically very similar organoids. This result suggest that provided a permissive microenvironment, the intrinsic genetic properties of cells are the one dictating architectural fate in 3D.

Together with its human origin, tissue-matched mechanical properties, balanced biochemical composition, and reduced pro-inflammatory bias, atdECM represents a robust, translationally relevant alternative to Matrigel. Beyond pancreatic organoids, the accessibility of human adipose tissue positions atdECM as a scalable and ethically favourable platform for advanced organoid, tumoroid, and disease-modeling applications, with the additional benefit of reducing reliance on animal-derived matrices.

## FIGURE LEGENDS

**Supplementary Figure 1: atdECM and Matrigel soluble proteins characterization**

(A) Cytokine expression in atdECM (n=2 donors and 3 membranes / donor) (Proteome profiler Human XL Cytokine Array Kit and Proteome Profiler Human Receptor Array, Non-Hematopoietic) and (B) in Matrigel (n=3) (Proteome Profiler Mouse XL cytokine Array Kit).

**Supplementary Figure 2: Proteomic dataset of atdECM samples and Matrigel obtained by mass spectrometry**

Excel file containing the mass spectrometry dataset of proteins identified in atdECM samples and in Matrigel, including annotated gene names, matrisome classification, matrisome category, Gene Ontology terms, molecular weight, number of identified peptides, sequence coverage and mean iBAQ values. The dashed line represents the median of all values.

**Supplementary Figure 3: Differential gene expression analysis between atdECM and Matrigel**

Excel file containing the results of DEG analysis between atdECM and uncoated condition performed using DESeq2. The table includes gene names, base Mean expression values, log2FoldChange, log2FoldChange standard error (lfcSE), test statistic (Wald test), p-values, adjusted p-values (padj), as well as the threshold and score used for DEG selection. Genes were considered differentially expressed with an adjusted p-value < 0.01 and log2FC > 1.

**Supplementary Figure 4: Differential gene expression analysis between atdECM and uncoated condition**

Excel file containing the results of DEG analysis between atdECM and uncoated condition performed using DESeq2. The table includes gene names, base Mean expression values, log2FoldChange, log2FoldChange standard error (lfcSE), test statistic (Wald test), p-values, adjusted p-values (padj), as well as the threshold and score used for DEG selection. Genes were considered differentially expressed with an adjusted p-value < 0.01 and log2FC > 1.

**Supplementary Figure 5: Differential gene expression analysis between Matrigel and uncoated condition**

Excel file containing the results of DEG analysis between atdECM and uncoated condition performed using DESeq2. The table includes gene names, base Mean expression values, log2FoldChange, log2FoldChange standard error (lfcSE), test statistic (Wald test), p-values, adjusted p-values (padj), as well as the threshold and score used for DEG selection. Genes were considered differentially expressed with an adjusted p-value < 0.01 and log2FC > 1.

**Supplementary Figure 6: Leading-edge gene from GSEA analysis**

Excel file containing the leading-edge genes identified by gene set enrichment analysis (GSEA). The table included the pathway name, normalized enrichment score (NES), adjusted p-value for the pathway (padj pathway), direction of regulation (up/down), gene name, log2FoldChange and adjusted p-value for each gene (padj gene).

**Supplementary Figure 7: Summary of enriched pathways from GSEA analysis**

Excel file summarizing the pathways identified as significantly enriched by gene set enrichment analysis (GSEA). The table included the pathway name, normalized enrichment score (NES), adjusted p-value for the pathway (padj pathway), direction of regulation (up/down), gene name, log2FoldChange and adjusted p-value for each gene (padj gene).

## Funding sources

FJ, TSB and CB were supported by the KATY project, which has received funding from the European Union’s Horizon 2020 research and innovation program under grant agreement No 101017453. FJ and CB was supported by the CANVAS project, which has received funding from the Horizon Europe twinning program under grant agreement No 101079510, and by the DIGPHAT project (Multi-scale and longitudinal data modeling in pharmacology: toward digital pharmacological twins), which has received funding from the French research initiative “France 2030” through the program PEPR Digital Health under ANR grant agreement No 22-PESN-0017. Some computations were performed using the GRICAD infrastructure (https://gricad.univ-grenoble-alpes.fr), which is supported by Grenoble research communities. LM is supported by the French National Research Agency in the framework of the ANR JCJC ADOC (ANR-24-CE19-3753-01).

This work was supported by the internal funding “Organoids on chip” FOCUS program of the CEA, the French Alternative Energies and Atomic Energy Commission and by the internal funding CFR Amont-Aval program of the CEA.

The proteomic experiments were partially supported by Agence Nationale de la Recherche under projects ProFI (Proteomics French Infrastructure, ANR-10-INBS-08 & ANR-24-INBS-0015) and GRAL, a program from the Chemistry Biology Health (CBH) Graduate School of University Grenoble Alpes (ANR-17-EURE-0003).

Sequencing was performed by the GenomEast platform, a member of the ‘France Genomique’ consortium (ANR-10-INBS-0009).

The microscopy facility MuLife of IRIG/DBSCI, funded by CEA Nanobio and GRAL LabEX (ANR-10-LABX-49-01) financed within the University Grenoble Alpes graduate school CBH-EUR-GS (ANR-17-EURE-0003).

## Acknowledgements

We thank also Denis Mariolle (CEA-Leti, Univ. Grenoble Alpes, F-38000 Grenoble, France) for its work on AFM data. Part of this work, carried out on the Platform for Nanocharacterisation (PFNC), was supported by the ‘Recherches Technologiques de Base’ program.

This article underwent linguistic refinement via *Maïa-Chat*, an AI tool developed by the French Alternative Energies and Atomic Energy Commission (CEA), to optimize clarity and precision.

