## Supplementary figures and images for "Human decellularized extracellular matrix from adipose tissue is a permissive microenvironment for pancreatic organoids generation"

### Supplementary Figure 2

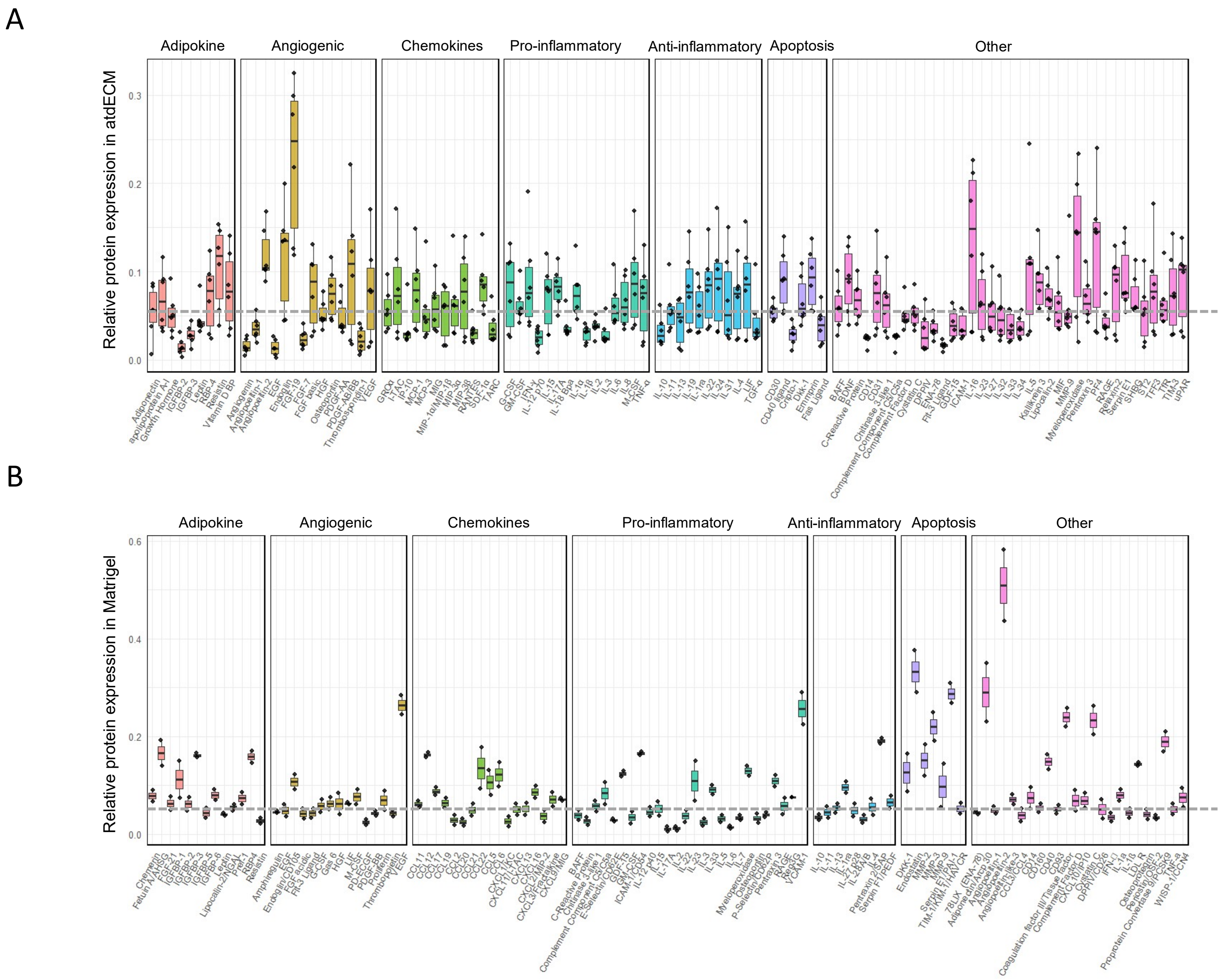
